# PAC: Highly accurate quantification of allelic gene expression for population and disease genetics

**DOI:** 10.1101/2021.07.13.452202

**Authors:** Anna Saukkonen, Helena Kilpinen, Alan Hodgkinson

## Abstract

Analysis of allele-specific gene expression (ASE) is a powerful approach for studying gene regulation. However, detection of ASE events relies on accurate alignment of RNA-sequencing reads, where challenges still remain. We have developed PAC, a method that combines multiple steps to improve the quantification of allelic reads, including personalised (i.e. diploid) read alignment with improved allocation of multi-mapping reads. We show that PAC outperforms standard alignment approaches for ASE detection in both accuracy and in the number of sites it can reliably quantify.

---

Allele-specific expression (ASE) is the imbalanced expression of the two alleles of a gene. While many genes are expressed equally from both alleles, gene regulatory differences driven by genetic changes (i.e. regulatory variants) frequently cause the two alleles to be expressed at different levels, resulting in allele-specific expression patterns. With RNA-sequencing (RNA-seq) data, the expression from the two alleles can be distinguished and quantified, but this analysis remains susceptible to many technical challenges, despite improved analytical methods^1,2^. To date, ASE analysis has largely been performed in the context of expression quantitative trait loci (eQTL) studies, as an alternative method to identify and characterise effects of regulatory variants on gene expression ^3,4^. In these studies, large sample sizes help mitigate the effects of technical biases^5^. However, the power of ASE analysis lies in its applicability to individual samples, particularly in the context of rare diseases and other cases where looking at combined haplotypic effects of multiple variants is necessary. Numerous studies have now leveraged the power of ASE in detecting genetic effects on gene regulation^6,7^ and identifying regulatory dysfunction in rare disease samples^8,9^. In order to draw conclusions about individual samples and loci, the accuracy of ASE calling becomes paramount.

One of the main sources of bias in ASE analysis is the alignment of sequencing reads. When short reads are aligned to the reference genome, reads carrying alleles that match to the reference sequence frequently map better than those carrying alternative alleles, leading to false ASE effects (‘reference allele bias’). Similarly, other factors such as reads that align to multiple locations in the genome can also influence the accuracy of allele counts at heterozygous sites. In the absence of better approaches, such difficult to map reads are typically discarded from the analysis^1^. Diploid genome mapping (i.e. use of personalised genome references) has been proposed as a solution to alignment-related artifacts in ASE analysis^10,11^, but these approaches do not deal with all potential causes of bias and they have not been widely adopted, likely due to the lack of a comprehensive pipeline to handle personalised genome coordinates in downstream analysis. Here, we address some of the remaining challenges in ASE analysis and describe the ‘Personalised ASE Caller’ (PAC) pipeline that integrates a series of existing and novel tools to improve the detection of genuine ASE events in short read RNA-seq data.

PAC implements a series of steps to align RNA-seq reads to a diploid reference genome and therefore more accurately detects and quantifies ASE events (**Figure 1A**). First, PAC uses phASER^12^ with read aware mode to improve variant phasing, particularly for rare variants, before creating parental genomes for the individual with the AlleleSeq software^10^. RNAseq reads are aligned to each genome individually with STAR^11^, retaining only properly paired and uniquely mapped reads, followed by a re-alignment step with RSEM^13^, which allows for the retention of multi-mapped reads through the implementation of an expectation-maximisation algorithm. For each pair of reads, PAC then selects the best location across the two parental genomes with custom scripts, before site-level and gene-level (phASER) allele counts are extracted genome-wide (see **Methods**). Throughout the development of PAC, we tested the effects of different read trimming and alignment parameters before arriving at the final optimised pipeline (see **Supplementary Materials**).

**Figure 1.**
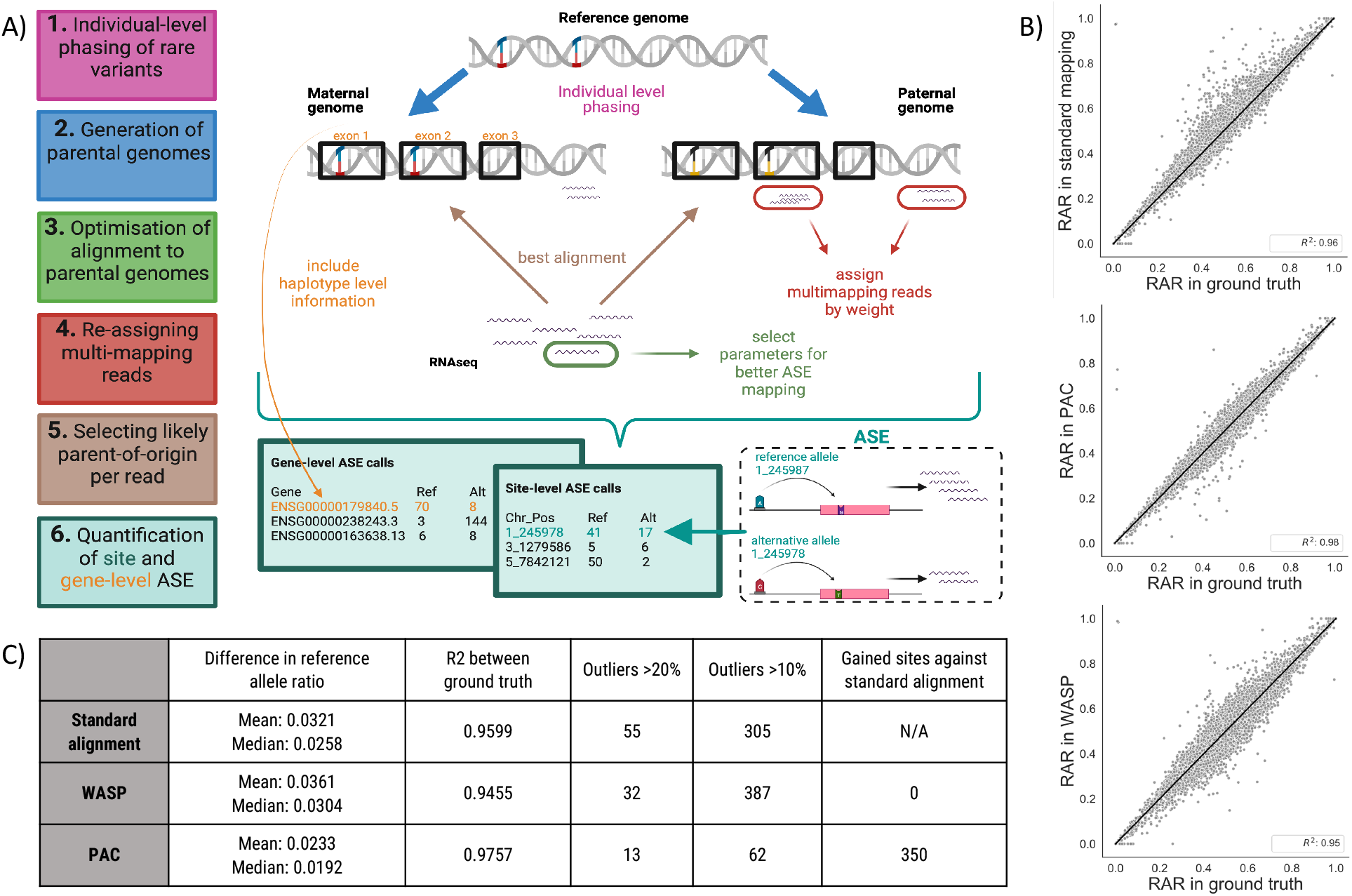
Overview of the PAC pipeline. **(a)** A schematic describing the main steps, features, and outputs of PAC. **(b)** Correlation of reference allele ratios (RAR) between the three different methods (standard alignment, WASP-filtered alignment, PAC) and the ground truth data. Genome-wide Pearson correlation coefficients (R^2^) are shown (P<0.05 for all comparisons). **(c)** Site-level summary statistics for the different analysis methods. Statistics are reported for sites with at least 20X coverage in all three methods.

To compare the performance of PAC with other approaches to quantify allelic expression, we generated simulated RNA-seq data where we knew the exact allele counts at each heterozygous site of a single individual (NA12877). To do this, we used high-quality genotype data from the Platinum Genomes Project^14^. First, whole-genome sequencing data were simulated using these variants, which was subsequently aligned to the reference genome before variant calling was performed with GATK^15^, followed by phasing with SHAPEIT2^16^ (see **Materials and Methods**). In this way, we generated a realistic set of variant calls that could be used for ASE analysis. Second, we generated simulated RNA-seq data for the individual using RSEM. To do this, we simulated RNA-seq data for each parental haplotype inherited by the offspring, before merging and calculating the ‘ground truth’ allele count for each allele at each heterozygous variant position identified in the GATK output (**Suppl. Figure 1**). This ground truth data was used as the baseline to which the accuracy of ASE calls obtained using different parameters and methods was compared. In total, the simulated RNA-seq data had a coverage of at least 20X at 13,211 unique heterozygous sites, 1,359 of which (10.3%) showed ASE under a standard binomial test (P < 0.05, corrected for 13,211 tests, **Suppl. Figure 2**).

Simulated RNA-seq reads were then aligned to the reference genome using three approaches: 1) standard alignment using STAR (to obtain a baseline set of ASE calls when using one of the most commonly used haploid alignment methods), 2) PAC (diploid alignment) and 3) WASP ^1^, a commonly used tool to correct for reference bias in ASE analysis. WASP incorporates a number of features to identify sites under ASE and whilst we cannot directly compare these with the other approaches, we can test the impact of read removal within the WASP pipeline. After applying each of the three approaches, we counted reads containing the two alleles at each heterozygous site and compared these data to the ground truth (**Figure 1C**). We focused on sites that had at least 20X coverage in both the ground truth data and all three alignment approaches (11,602 heterozygous sites) to make results directly comparable (**Figure 1B**).

First, with standard alignment, ~91.7% of sites that had at least 20X coverage in the ground truth data also met this coverage threshold (12,109/13,211), with the average coverage at these sites dropping from ~175X in ground truth data to ~144X with standard alignment, highlighting the loss of many reads when using the standard approach. We find that reference allele ratios (RAR) showed a high correlation between standard alignment and the ground truth data at heterozygous sites (R^2^=0.960) (**Figure 1C**). However, 305 sites show an absolute difference in RAR of greater than 10% and 55 sites a difference greater than 20%, The absolute mean difference shows a 3.21% bias across all heterozygous sites (**Figure 1B**).

Second, when we applied the PAC pipeline to the simulated data we find that the number of reads and the accuracy of allelic assignment is significantly improved compared to standard alignment. First, the number of sites that have at least 20X coverage in both ground truth and PAC-aligned data increased to 12,448 (339 additional sites compared to standard alignment), with an average coverage of ~150X at these sites. Second, the correlation of RAR in PAC compared to the ground truth data increased to R^2^=0.976 (**Figure 1C**). The number of outlier sites also dramatically decreased to 62 and 13 sites showing an absolute difference in reference allele ratio of >10% and >20%, respectively. The mean difference from ground truth RAR is 2.33% (**Figure 1B**), which is significantly lower than that found for standard alignment at the same sites (one-sided t-test, P=2.6×10^−125^).

Third, we compared WASP-filtered^1^ data to the ground truth and found that the number of sites that have at least 20X coverage in both ground truth and WASP-corrected data dropped to 11,612 (836 fewer than when using PAC), with an average coverage of 135X. Both of these results are likely a consequence of the approach used in WASP to remove difficult-to-align reads from the analysis. While the number of extreme outliers reduced to 32 (absolute difference of >20%) using WASP-filtered data, the number of sites with an absolute difference in RAR of >10% increased to 387 compared to standard alignment, and the R^2^ value decreased to 0.946 (**Figure 1C**). Furthermore, the mean absolute difference between WASP-filtered data and the ground truth (3.61%) is significantly higher than both standard alignment (P=4.5×10^−21^, one-sided t-test) and PAC (P=8.9×10^−272^, one-sided t-test) (**Figure 1B**).

Compared to both standard and WASP-filtered alignment, applying PAC results in an additional 350 heterozygous sites that have coverage of at least 20X that do not meet this threshold in both of the other two approaches. Allele count quantification at these sites is also highly accurate with PAC (R^2^=0.844, compared to the ground truth). Similarly, for sites detected in standard alignment and PAC (but not WASP-filtered data), PAC quantification is highly significant (496 sites, R^2^=0.956, P=2.6×10^−266^, **Figure 2A**), showing that PAC performs well at sites with lower coverage that may be missed by other approaches. At least part of the improvement in accuracy when using PAC appears to occur in regions of the genome where accurate alignment is known to be more difficult. For example, the difference in RAR from the ground truth is significantly higher in standard aligment and WASP-filtered data than PAC at sites where there is an indel (>6bp) within 500bp of the heterozygous site (**Figure 2B**). The trend is similar when another heterozygous site or a rare variant (MAF <1%) is close by (**Figure 2B**).

**Figure 2.**
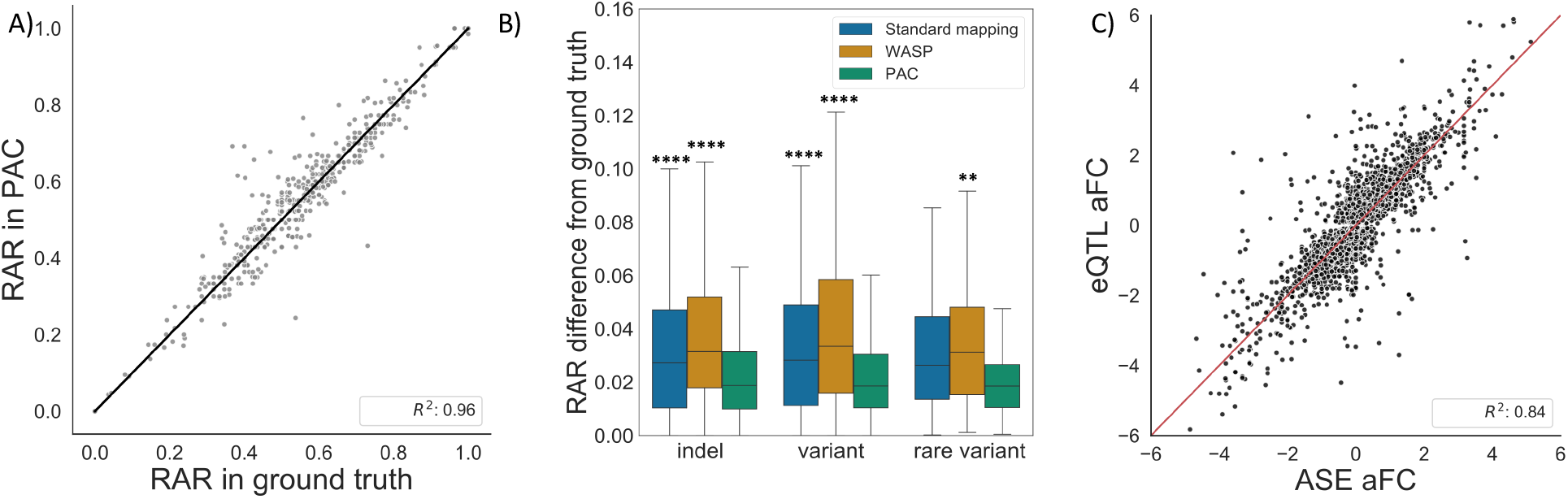
Performance of PAC compared to other methods. **(a)** Genome-wide correlation of reference allele ratios (RAR) at heterozygous sites that PAC and standard alignment detects but that are discarded by WASP-filtering (Pearson correlation R^2^=0.956, P=2.6×10^−266^). Sites with at least 20X coverage were considered. **(b)** The difference in reference allele ratio of sites that are within 500bp of at least 6bp indel, within 25bp of another variant or a rare (MAF <1%) variant in different analyses against the ground truth. Sites shared between all methods and with at least 20X coverage were considered. Mann-Whitney test was performed with Bonferroni correction to adjust for multiple testing (P<=1e-04 for ****; 1.00e-03 < P <= for 1.00e-02**) and stars above each boxplot refer to the comparison against PAC **(c)** Correlation of allelic fold change (aFC) values derived from ASE and eQTL analyses from 670 GTEx whole blood samples. Genes with a significant eQTL (q-value < 5%) and gene-level ASE information for at least 10 individuals were selected. Pearson correlation coefficients are shown for eQTL versus ASE aFCs derived using PAC (see also **Supplementary Figure 3**).

Finally, to characterise the performance of PAC on population level data, we aligned 670 whole blood RNA-seq samples from the GTEx project (v8)^4^ using PAC, obtained gene-level counts for each gene and individual, and then compared the allelic fold change (aFC)^17^ for each gene generated from the count data with those obtained from eQTL mapping. We also compared these results to those generated through standard and WASP-filtered alignments using gene-level count data from Castel *et al*^18^. Using genes with a significant eQTL (q-value < 5%) and where the aFC could be calculated from ASE data in all three methods (8913 genes, see **Materials and Methods**), PAC shows the strongest correlation between ASE and eQTL aFC (R^2^=0.842, **Figure 2C**), followed by WASP-filtered data (R^2^=0.829) and then standard alignment (R^2^=0.820). Furthermore, due to the higher coverage obtained when aligning data with PAC, we were able to generate aFC for an additional 740 genes using this approach that did not meet coverage criteria in WASP-filtered data, and the aFC generated from ASE and eQTL data was still highly correlated amongst these genes (R^2^=0.653, P=4.0×10^−91^). Similarly, there were 319 genes present in PAC data that were not in either standard or WASP-filtered alignments; again the correlation between ASE and eQTL aFC for these genes was significant (R^2^=0.643, P=1.1×10^−38^). These results indicate that PAC not only has improved accuracy, but also more power to detect the influence of regulatory variation on gene expression across a larger number of genes.

In summary, we present PAC, a new tool for ASE analysis that generates highly accurate allele counts from RNA-seq data for use in studying the regulation of gene expression. PAC is written in Nextflow and is available to use via https://github.com/anna-saukkonen/PAC. Our approach incorporates both novel and existing analytical steps to better control for common technical biases that occur when aligning short read data, including reference allele bias and reads that align to multiple different genomic locations. Additionally, PAC maximises ASE quantification accuracy for individual samples by improved phasing of rare variants and subsequent diploid genome alignment. Using both simulated and population level RNA-seq data, we show that PAC performs better than standard alignment techniques and other commonly used tools that attempt to deal with some of the technical issues related to ASE analysis, producing both more accurate allele counts and higher coverage at heterozygous sites. We anticipate PAC to be primarily useful in the context of rare diseases and other situations where small sample size precludes the use of population level methods to study differences in gene expression and regulation. In such cases, accurate quantification of allelic expression changes, in individual samples, is of paramount importance in understanding disease biology. However, PAC can also be used on population level data, where allelic imbalance information can be used to better infer the impact of genetic variants on the expression of nearby genes. In these ways, PAC can be applied to the vast quantity of existing RNA-seq datasets to better understand a wide array of fundamental biological and disease processes.

## Materials and Methods

The materials and methods are described in full in the **Supplementary Material.**

## Acknowledgements

AS is supported by the London Interdisciplinary (LIDO) Doctoral Programme funded by the UK Biotechnology and Biological Sciences Research Council (BBSRC). HK and AH have both received support from MRC eMedLab Medical Bioinformatics career development awards from the UK Medical Research Council (MR/L016311/1). This work is (partly) funded by the NIHR GOSH BRC. The views expressed are those of the author(s) and not necessarily those of the NHS, the NIHR or the Department of Health. The authors acknowledge use of the research computing facility at King’s College London, Rosalind (https://rosalind.kcl.ac.uk), which is delivered in partnership with the National Institute for Health Research (NIHR) Biomedical Research Centres at South London & Maudsley and Guy’s & St. Thomas’ NHS Foundation Trusts, and part-funded by capital equipment grants from the Maudsley Charity (award 980) and Guy’s & St. Thomas’ Charity (TR130505). The views expressed are those of the author(s) and not necessarily those of the NHS, the NIHR, King’s College London, or the Department of Health and Social Care. The Genotype-Tissue Expression (GTEx) Project was supported by the Common Fund of the Office of the Director of the National Institutes of Health, and by NCI, NHGRI, NHLBI, NIDA, NIMH, and NINDS. The data used for the analyses described in this manuscript were obtained from the GTEx Portal and dbGaP accession number phs000424.v8.p2.

## Supplementary Material

Supplementary Material is provided as a separate PDF which includes the following:

- **Supplementary Materials and Methods**
- **Supplementary Figures 1-3.**
- **Supplementary Table 1.**

## Data and code availability

The PAC pipeline is available at https://github.com/anna-saukkonen/PAC. All data and other files used in this work are available and documented on Github (https://github.com/anna-saukkonen/PAC/article_data).

