## Supplementary Material for "PAC: Highly accurate quantification of allelic gene expression for population and disease genetics"

This document contains the following supplementary material for the manuscript:

1. Materials and Methods
2. Supplementary Results
3. Supplementary Figures 1-3
4. Supplementary Table 1

### Materials and Methods

*Ground truth variant calls*

Ground truth variant data was obtained for a single individual from the Platinum Genome Project (PGP), specifically, NA12877 from CEPH/Utah pedigree 1463[1]. The PGP generated deep (50X average) whole-genome sequencing (WGS) data from 17 individuals in a three-generation pedigree. The project used two different sequencing technologies and variant calls from six different informatics pipelines. Conflicts between call sets were resolved using inheritance-based validation. This dataset is widely considered to represent the most accurate set of variant calls that can be achieved with current methods. We used phased variant calls (VCF file) for NA12877 that included indels and SNPs to generate a diploid genome reference using AlleleSeq[2] and hg19 version of the human reference genome.

*Simulated WGS data*

The maternal and paternal genomes from AlleleSeq were used to simulate whole genome sequencing reads for NA12877 with ART[3]. As input, ART requires parameters related to insert size, read length, coverage, and standard deviation of fragment length. To obtain a realistic simulation, we acquired these parameters from real sequencing data for sample HPSI0114i-eipl_1 from the HipSci project[4]. WGS reads for 'eipl_1' were aligned to the hg19 reference genome using BWA-MEM[5]. Samtools[6] was then used to limit the output to uniquely mapping, properly-paired reads, and to obtain the required parameters for ART (from *samtools-stats*). Parameters were set as follows: 1) 20X coverage, 2) read length 150bp, 3) mean fragment length 479bp, and 4) standard deviation of fragment size 117bp. The maternal and paternal reads were simulated separately and then merged so that the final coverage of the simulated WGS sample was 40X.

*Variant calling*

Simulated WGS reads were aligned to the reference genome (hg19) the same way as described above, and variant calling was performed with GATK v. 4.0.12.0 according to recommended best practises[7]. Variants were then phased with Shapeit2[8] using the 1000 Genomes phase 3 reference panel. These variants were then compared against the ground truth variants from PGP (see **Supplementary Results**).

*Simulation of RNA-sequencing data*

In order to construct ground truth allele counts for NA12877, we simulated RNA-sequencing (RNA-seq) reads using RSEM v1.3.1[9]. To obtain realistic input parameters for RSEM, we used real data from the parents of this individual (NA12889 and NA12890, **Supplementary Figure 1**). Raw RNA-seq data for the parents (from lymphoblastoid cell lines) was obtained from the Geuvadis Project[10], trimmed, and mapped to the hg19 reference genome using STAR v.2.5.1a[11] with default parameters. The mapped reads from the parents (two separate bam files) were then input into RSEM to generate a single matrix of expression levels for each transcript in the Gencode v19 annotations. To simulate RNA-seq reads, *rsem-simulate-reads* function was used with the following input parameters obtained from the alignment of data from NA12889 and NA12890: (i) fraction of reads coming from background noise (0.27 and 0.19, paternal and maternal sample, respectively) and (ii) total number of reads to be simulated (40.9M and 28.1M paternal and maternal sample, respectively). Since the input for the simulations is based on real data from two distinct individuals, this generates allelic variation at heterozygous positions of the genome, i.e. ASE effects in the simulated data. To acquire ‘ground truth allele counts’, we obtained maternal and paternal allele counts at heterozygous genome positions of NA12877 from the PGP VCF file. The simulated reads from the parents (FASTA files) were then merged into a single RNA-seq sample, representing the (simulated) transcriptome of individual NA12877.

*Ground truth allele counts*

The genomic coordinates of the simulated RNA-seq reads were obtained using custom scripts, based on the flag information stored for each read by RSEM (including transcript ID, position on transcript, etc.). LiftOver was used to convert read locations from the parental genomes to the reference genome using chain files generated by AlleleSeq. We then counted the number of reads from reference and alternative alleles that overlapped all heterozygous positions in NA12877 based on the PGP variant calls. These allele counts were then combined for each site to create ground truth allele counts in the offspring. For all subsequent analyses, we used heterozygous sites with at least 20X read coverage (sum of reference and alternative allele counts). The distribution of the reference allele ratios across all 20X sites in the ground truth data is shown in **Supplementary Figure 2**.

*Standard alignment of simulated RNA-seq reads*

The paired-end reads were aligned to a reference genome (1000G version of GRCh37) with STAR 2.51a with standard parameters including soft-clipping, using two-pass mapping, version 19 of the Gencode gene annotation and allowing 8 mismatches per read pair, before keeping only properly paired (-f 0x0002 using Samtools) and uniquely mapped (NH:i:1 flag) reads.

*PAC pipeline*

The final PAC pipeline (**Figure 1**) was constructed as follows, with each step tested for its impact on the accurate alignment of reads compared to the ground truth data (see **Supplementary Table 1**). First, standard alignment of RNA-seq reads was performed (see above). These data were then used as input for phASER[12], alongside the phased VCF obtained from the GATK pipeline above, to re-determine the phase at heterozygous sites where RNA-seq reads can add information (read aware mode). The resulting VCF file was then used with AlleleSeq[2] to generate parental genomes. For each parental genome, simulated RNA-seq reads were aligned with STAR as above, keeping only properly paired and uniquely aligned reads. Since the uniquely alignable regions in the reference and non-reference genome may differ, we also used RSEM (v1.3.0)[9] to take the original alignment from STAR (containing all reads aligned to transcriptome coordinates, including reads that align to multiple locations) and re-align the data using the --sampling-for-bam flag to output a single location for each read based on its posterior probability generated from estimated abundances. Additional reads aligned by RSEM that were not uniquely aligned using STAR were then added to the final bam file. After alignment of each parental genome, a custom script was used to select the best alignment for each read from the two mappings (scoring reads by the number of matching nucleotides minus two times the number of indel positions, drawing at random when the two mappings have equal scores), and the number of each allele at each heterozygous site was counted. We also produce allele counts at haplotypic level using phASER Gene AE. PAC is available at https://github.com/anna-saukkonen/PAC along with full instructions of use.

*WASP*

WASP-filtering was performed using the same approach as detailed above for standard alignment, but using the additional flag --waspOutputMode SAMtag within STAR (v2.7.3a), together with providing the VCF file generated from GATK (as described above). We filtered the resulting BAM file for reads that were properly paired, reads without a WASP flag (and thus do not contain a genetic variant) and reads that pass WASP-filtering (with flag 'vW:i:1'), before counting reference and alternative alleles at heterozygous sites

*Evaluation of allele count accuracy / Outlier analysis*

In order to evaluate the performance of the pipeline and how the different steps influence the accuracy of allele counts (and eventual ASE calls), we compared results obtained with PAC, standard alignment and WASP-filtering to the ground truth data. We excluded sites that were located in the HLA region as well as the blacklisted genomic regions. We monitored the number of 'accessible' heterozygous sites (i.e. bi-allelic sites that had at least 20X coverage, obtained through *samtools mpileup* using default parameters and disabling read-pair overlap detection), correlation of the reference allele ratios (RAR) in the analysis and the ground truth, sites that were present in standard alignment but missed by analysis, and the number of sites where the RAR showed more than 10% or 20% difference between the analysis and the ground truth (referred to as outliers).

*Accuracy of analysis near indels and other variants*

To evaluate how PAC performs at genomic regions that are difficult to align relative to standard alignment and WASP-filtered data, we compared the difference in RAR against ground truth RAR at heterozygous sites. For indel analysis, we selected sites that were within 500bp of an indel (minimum indel length 6bp). We also looked at sites that had another heterozygous single nucleotide variant or rare variant (MAF < 1%) within 25bp of the heterozygous site. We used CEU population data from 1000 genomes project for this. Mann-Whitney test was performed with Bonferroni correction to adjust for multiple testing (**Figure 2B**).

*Analysis of GTEx samples*

To interrogate the performance of PAC on population level data, we obtained aligned data containing all reads for 670 whole blood samples from the GTEx project (v8, aligned to the hg38 reference genome)[13], converted these files back to raw sequence files (fastq) with Samtools, and then used them as input for PAC (selecting the GRCh38 reference genome), together with phased genetic variant calls from WGS data (obtained from the GTEx, phASER_GTEx_v8_merged.vcf.gz). Gene level count data was then obtained from the output of PAC and used to calculate allelic fold change (aFC) estimates per gene using pHASER-POP[14], which retains only genes and samples with at least 8 read counts. Within phASER-POP, we supplied lead eQTL variants identified in the GTEx project (v8) for each gene, which we obtained from the GTEx portal (Whole_Blood.v8.egenes.txt.gz). We also ran phASER-POP using two additional gene count matrix files representing standard alignment and WASP-filtered alignment, both obtained through the GTEx portal (phASER_GTEx_v8_matrix.gw_phased.txt.gz and phASER_WASP_GTEx_v8_matrix.gw_phased.txt.gz, respectively) and produced by Castel *et al.*[14]. We then compared the aFC for each gene generated using ASE data where at least ten individuals were heterozygous for the lead eQTL variant linked to the gene, with aFC estimates generated from eQTL data after filtering genes where the eQTL association was q-value < 5%. We selected only genes that were present in all three methods for direct comparison.

**Supplementary Results**

*Assessment of variant calling accuracy on ASE detection*

Accuracy of variant calls is a known source of error in ASE analysis. For example, variants erroneously called as heterozygous would show complete monoallelic expression in ASE analysis. Further, while ASE analysis is typically carried out only with SNVs, the call accuracy of short indels and copy number variants (CNV) can influence the mapping of reads when personalised genomes are used. As such, we analysed the ability of GATK to identify heterozygous variants using simulated whole genome sequencing data as described in the methods. In total there are 4,042,773 data points in the PGP VCF file for NA12877 and 4,011,226 in the GATK output, showing a true positive rate of 99.22%. The GATK VCF file also contains an additional 5,389 data records not present in the original data, leading to a false positive rate of 0.134%. The GATK VCF misses 36,939 data points, with false negative rate being 0.914%.

Following this, we generated two parental genomes within AlleleSeq[2] using phased DNA variant calls from PGP for NA12877 and aligned simulated RNA-seq data to each genome individually before selecting the best alignment for each read pair for the two mappings (see **Materials and Methods**). In the ground truth data there are 13,211 heterozygous sites that have at least 20X coverage. Using this alignment approach, the number of heterozygous sites that also have at least 20X coverage was 12,405, with an average coverage of ~149X, and the correlation between the reference allele ratio at these sites for the aligned data versus the ground truth was $R^2$=0.962. In total, 161 sites showed an absolute difference in reference allele ratio (versus the ground truth) of greater than 10%, and 72 sites showed a ratio >20% (**Supplementary Table 1**).

*Effects of read trimming and softclipping*

Many RNA-seq processing pipelines perform read trimming for adaptors, base quality and polyA tails before alignment. In ASE analysis, this may lead to biased results if alternative alleles at the ends of reads are preferentially trimmed. As such, we compared the effects of read trimming against using soft-clipping within STAR on allele counts at heterozygous sites, and found a much closer agreement with the ground truth for the latter approach. Including read trimming (stringency of 3bp, removing adaptors and terminal bases with phred qualities lower than 30), the number of heterozygous sites with at least 20X coverage in both aligned and ground truth data was 12,161 and the correlation of reference allele ratios versus the ground truth decreased to $R^2$=0.947. Similarly the number of sites showing an absolute difference in reference allele ratio of >10% and >20% was 252 and 69, respectively (**Supplementary Table 1**), showing that applying softclipping with STAR is favourable to read trimming in these circumstances.

*Effects of local phasing of variants (phASER)*

Accurate phasing of genetic variants impacts the ability to construct correct parental genomes. Similarly, it is difficult to accurately phase rare alleles if they are not present in reference datasets. As such, we used the read aware mode of phASER [12] within our pipeline, which improves local phasing by considering whether nearby genetic variants fall on the same or opposite reads (or pairs). Using this approach, we see a marginal improvement in all parameters: 12,415 heterozygous sites have coverage of at least 20X in aligned data, the correlation between reference allele ratios increases slightly to $R^2$=0.963 (versus the ground truth), and the number of sites showing an absolute difference in reference allele ratio between ground truth and aligned data decreases slightly to 157 for differences >10% and 68 for differences >20% (**Supplementary Table 1**).

*Effects of recovering multi-mapping reads*

In many RNA-seq experiments, reads that do not align uniquely to the reference genome are typically discarded. For ASE analysis, this can lead to biased allele counts at specific loci, if the expression level of one or both alleles is underestimated due to non-unique alignment. Methods to recover reads that align to multiple locations (from hereon ‘multi-mapping’ reads) exist, but current ASE detection methods do not incorporate these reads into the analysis. In order to include such reads, we used RSEM within our pipeline to assign a single location for multi-mapping reads based on read depth of uniquely aligned reads (see **Materials and Methods**) for each parental genome, and then incorporated these reads into the final aligned files before selecting the best alignment for each read pair for the two mappings. Applying this approach, we again improve the accuracy of the alignment, with 12,448 heterozygous sites having at least 20X coverage, the correlation of reference allele proportion at the sites between the ground truth and aligned data of $R^2$=0.968, and 140 and 46 sites showing an absolute difference in reference allele ratio of >10% and >20%, respectively (**Supplementary Table 1**).

**Supplementary Figures**

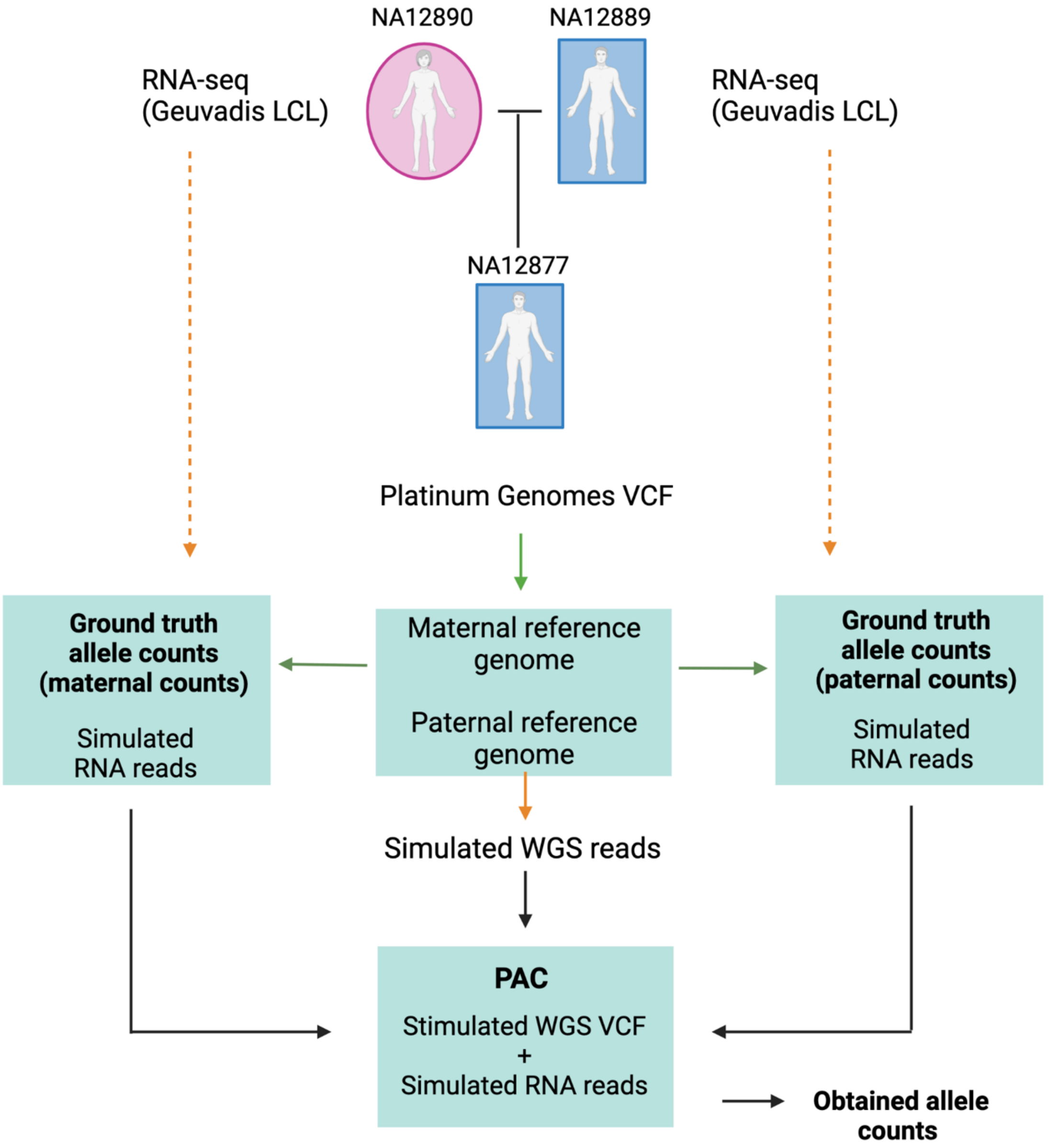


**Supplementary Figure 1. Ground truth data generation for individual NA12877.** In Platinum Genomes VCF, variants were verified using multiple sequencing platforms and analysis methods, and conflicting calls resolved using parental genomic information. Platinum Genomes VCF together with a reference genome were used to generate ground truth genomes using AlleleSeq. Ground truth genomes were used to simulate RNA sequencing reads using RSEM. Parameters were obtained from RNA-sequencing reads of LCLs from individuals NA12890 and 12889 that were the actual parents, obtained from the Geuvadis Project. The simulated RNA reads were used to count coverage at each heterozygous site, called ground truth allele counts. Ground truth genomes were also used to simulate WGS using ART. Parameters for this were obtained from HipSci sample HPSI0114i-eipl_1. Simulated WGS were used to obtain variant calls using GATK best practises. This VCF, together with simulated RNA-seq reads, were used for PAC to obtain allelic count data that were compared against ground truth allele counts at heterozygous sites.

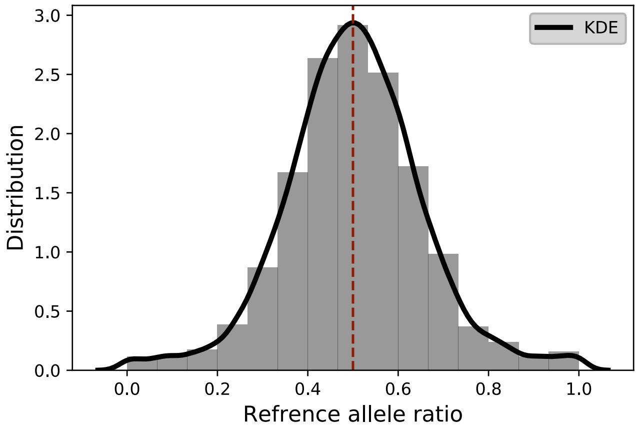


**Supplementary Figure 2. Distribution of reference allele ratios in ground truth data.** All heterozygous sites in ground truth data with 20X coverage are shown. Ratio of 0.5 implies that both alleles are expressed at equal ratios. KDE = kernel density estimate.

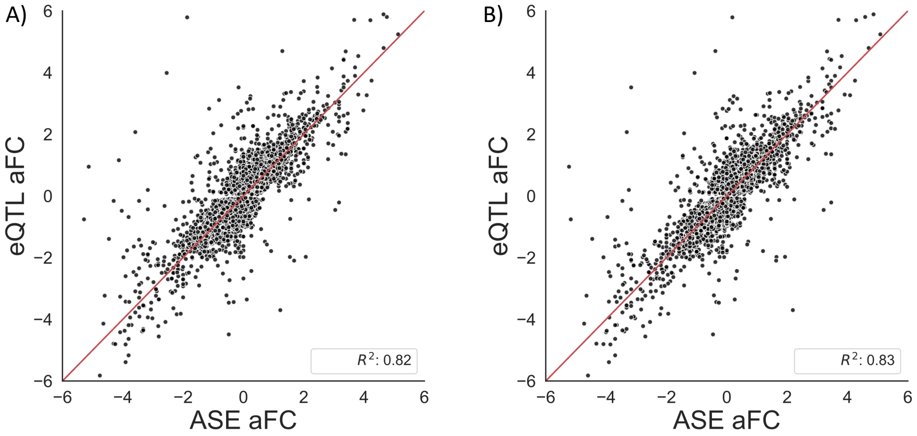


**Supplementary Figure 3. Gene-wise comparison of ASE and eQTL signals.** Correlation of allelic fold change (aFC) values derived from ASE and eQTL analyses from 670 GTEx whole blood samples. Genes with a significant eQTL (q-value < 5%) and gene-level ASE information for at least 10 individuals were selected. Pearson correlation coefficients are shown for eQTL versus ASE aFCs derived using standard alignment **(a)** and WASP **(b)**.

**Supplementary Table 1. Summary of PAC parameter optimisation.** Multiple different steps were tested for their impact of allelic read counts, including trimming of adaptors and low quality nucleotides (TRIM), soft-clipping in STAR (SOFT), read-aware phasing in pHASER (PHASE) and reallocation of multi-mapped reads (MULTIMAP) (see **methods**).

| | TRIM PHASE NO MULTIMAP | TRIM PHASE MULTIMAP | SOFT PHASE MULTIMAP | SOFT PHASE NO MULTIMAP | TRIM NO PHASE NO MULTIMAP | SOFT NO PHASE MULTIMAP | SOFT NO PHASE MULTIMAP | SOFT NO PHASE NO MULTIMAP |
|---|---|---|---|---|---|---|---|---|
| Sites shared with ground truth | 12159 | 12194 | 12448 | 12415 | 12161 | 12190 | 12436 | 12405 |
| Difference in reference allele ratio | Mean: 0.0331 Median: 0.0273 | Mean: 0.0326 Median: 0.0273 | Mean: 0.0248 Median: 0.0195 | Mean: 0.0254 Median: 0.0196 | Mean: 0.0332 Median: 0.0273 | Mean: 0.0326 Median: 0.0272 | Mean: 0.0249 Median: 0.0196 | Mean: 0.0255 Median: 0.0196 |
| R2 between ground truth | 0.9475 | 0.9532 | 0.9681 | 0.96251 | 0.9469 | 0.9530 | 0.9679 | 0.9619 |
| Outliers >20% | 67 | 47 | 46 | 68 | 69 | 49 | 49 | 72 |
| Outliers >10% | 246 | 240 | 140 | 157 | 252 | 244 | 144 | 161 |
| Sites not in standard alignment | 209 | 242 | 350 | 318 | 207 | 233 | 339 | 311 |
| Sites not in WASP | 606 | 640 | 846 | 813 | 605 | 632 | 833 | 802 |
